# Matriarchy and prehistory: a statistical method for testing an old theory

**DOI:** 10.1101/141374

**Authors:** Julien d’Huy

## Abstract

Mythological data and statistical methods have been used to reconstruct the phylogeny of matriarchal tales and its relationship to genetic trees. The results show a correlation between the two trees and allow to identify the proto-forms of this mythology. This makes it possible to reject some assumptions about rock art.

## 1. Introduction

The hypothesis of the matriarchy as a social model, based on the maternal rights and hegemony of women, finds its origin in the work of Johann Jakob Bachofen, *Das Mutterrecht* (1861). His work is chiefly based on the Greek myths, and the historical data left by earlier authors such Herodote, and follows the earlier *Mœurs des sauvages Américains comparées aux mœurs des premiers hommes* of Joseph François Lafitau (1724) which compares the supposed gynecocracy of the Hurons and Iroquois with that of the ancient Greeks and Egyptians. Johann Jakob Bachofen defends the existence of a matriarchal society in Ancient Greece: he evokes a primitive domination of the feminine principle in the religion and insists on the importance of the matrilineal filiation, in which the heredity of power is only transmitted from mother to girl. This matriarchal state would have preceded the patrilineal filiation, and would have followed a generalised stage of reproduction governed by the *ius naturale* of pleasure, where only maternity could be proved. And, in reaction to the abuses of men, the resentment of women leads to a gynecocracy and sometimes even to a community of Amazons. The men would then have to return to hegemonic power, changing the cult of the Earth-mother to a cult of the celestial gods. Bachofen’s work has influenced in a significant way many authors like John Lubbock (1867: 67–70), Lewis H. Morgan (1877), Friedrich Engels (1900) and Sigmund Freud (1922: 193). Many other authors, such as Jean Przyluski (1950) and Wilhelm Schmidt (1955), have contributed to popularising this matriarchal view. The first application of the hypothesis to rock art, if I am not mistaken, goes back to 1931, when Piotr P. Efimenko interpreted the feminine representations of the Upper Palaeolithic as a reflection of the social and economic role of the women at this time, as well as her spiritual role within a social organisation of a matriarchal type, dominated by the mothers.

And yet, outside these myths, matriarchy may never have existed. No example of gynecocracy has ever been observed in the world. Matriarchy or gynecocracy, where power is exercised exclusively or mainly by women, is a situation for which no clear examples have ever been found^1^ (Barry *and al.* 2000: 728; Le Quellec 2009: 247–250). As François Héritier (2011) has said: “societies where the power would be in the hands of the women with dominated men do not exist and never have existed. (…) There are no matriarchal societies, because the archaic model dominating all the planet comes from the beginning.^2^” If this narrative of a primitive matriarchy spread through all the planet it may have been at the same time as the first human migrations. The current matriarchal myths probably reflect not a previous state of human civilisation, but only a prehistoric myth.

Following this trail, Eugène Beauvois (1904: 324–326) incorrectly proposed that the Amazon-story must have reached America across the Atlantic Ocean through some Pre Columbian immigration of Celtic missionarie and Gudmund Hatt (1949: 72) noted that if one would seek a foreign origin for the American Amazon-stories, it should be in Asia and Oceania. According to Yuri Berezkin (2013: 142) “there are similar rituals/myths in Africa, Australia, Melanesia and South America that are associated with institutionalised opposition of the sexes, and typically include stories about the past domination of women, not only in the social sphere (the story of “a woman’s realm” sometime in the past, or in a distant land, is a motif known all over the world), but also in cult and ritual^3^”. And Yuri Berezkin further states that “this worldwide correspondence between narratives and rites, except in Eurasia, implies that we can assume that the male rituals associated with institutionalised opposition originated in Africa and were brought to Australia and New Guinea by the first sapiens. From East Asia, this mythological complex entered the New World along the coast with the first wave of migrants. […] Then these myths and rituals were gradually supplanted, entirely in Asia and mainly in North America, by new incomers from the Eurasian cultural household^4^” (Berezkin 2013: 147–148). He notes that this hypothesis would be confirmed by the diffusion of the motif F8 (according to his classification, i.e. : Initially women and men live apart from each other. Later they meet each other and come to live together), F38 (Women were possessors of the sacred knowledge, sanctuaries or ritual objects which are now taboo for them or they made attempts to acquire such a knowledge or objects), and F40B (A man gets into the village of women. Usually he has to satisfy every woman against his will or every woman claims him for herself), which would be explained by an Out-of-Africa diffusion.

According to the American anthropologist Joan Bamberger, principal function of the myth of the primitive matriarchy is to relegate the power of the women to a very remote past in order to guarantee the present state of the domination of one sex over the other^5^ (Bamberger, 1974: 280). But if Yuri Berezkin is right, the matriarchal myth already existed during the Out-of-Africa period. That implies that this myth could pass to the later cultures, ant that matriachy never existed. How can this be statistically demonstrated?

## 2. Material and method

The first to have applied the phylogenetic method to the mythology is, to my knowledge, Thomas Abler who in 1987 used the techniques of cluster analysis to classify and show relationships among the 41 variants from the Iroquoian myth of the creation of the world. He showed that most clusters of the trees represent a national or tribal tradition. The idea to use phylogenetic tools to classify tales or many versions of the same tale was taken up by Jun’ichi Oda (2001) and Jamie Tehrani (2013). Since 2012, I have used similar tools to trace lines of transmission of myths, folktales and mythological traditions back to a common ancestor and to test hypothesis about human prehistory and the process explaining the current geographic distribution of the myths and folktales, quantify patterns of borrowing, reconstructing proto-mythology and testing claims, such as the rate of evolution, about mythological evolution.

Myths are not genes: the biologist’s softwares have been used in this particular field of study for convenience. These software allow to construct trees based on similarities and differences between the studied units. If we accept that two people genealogically close share more narratives in common than two more distant people, or that their tales have more in common with each other than with more distant people, such softwares allow us to reconstruct “trees” of oral traditions (d’Huy 2012a, 2015b). However, these results mean nothing without a point of comparison, and need to be tested and confronted with other results.

I used the analytical catalog of the mythological motifs, graciously made accessible by Yuri Berezkin (http://ruthenia.ru/folklore/berezkin, consulted on 22/02/2016), to identify narratives about the differentiations between men and women, their cohabitation and their confrontation. Narratives as basic units to conduct phylogenetic studies have been adopted many times before (d’Huy 2015, Ross and Atkinson 2016, da Silva and Tehrani 2016). The Berezkin’s database is made up of 19th, 20th and 21st century simplified subjective summaries of longer secondary-source summaries of native myths in diverse genre of widely varying types and accuracy. It is necessary to note that such narratives support easily the translation from one language to another: a translation doesn’t affect the fund of tales, only their form. Moreover, Yuri Berezkin classifies the tales on an abstract level; the detail of the text, and its many alternate versions, are also not very important if the motif itself remains intact. Finally, numerous works, since Charles-Félix-Hyacinthe Gouhier in 1892, showed the durability through time of such motifs (also see, among others, Bogoras 1902, Jochelson 1905, Hatt 1949, Korotayev and Khaltourina 2011, Berezkin 2013, Witzel 2013 and Le Quellec 2014, 2015a, b). The continuous and repeated transformations and simplifications that take place in oral myths when they were passed into literate form have little impact on our analysis which is quantitative, on an abstract level, and not qualitative.

The cultural areas determined by Yuri Berezkin have also been considered as a unit of analysis. Such a choice implies that cultural areas are more than a convenient fiction: they are a real thing with ancestors, descendants and relations. Moreover, such a unit should be able to endure phenomena and persist over time. An advantage of this approach is that if a narrative has been lost in a given ethnic group, the neighbouring tribe of the same cultural area might remember it.

The following features were selected among the Berezkin’s corpus F1, F5, F5A, F8, F16, F16B, F16C, F16D, F38, F40A, F40B, F41, F42, F42A, F43, F43A, F43B, F43C, F44, F45, F45A, F45B, F46, F46A, F47, F47A, F48 and F97 (F39 being not available online). For every cultural area, each motif were coded by 1 if it was present, by 0 if it was absent. Then, to avoid any bias in the study, only geographical areas possessing at least eight narratives were retained. Finally, F45A that is not present in the Berezkin’s corpus has been deleted from the dataset.

Tools, usually used by the biologists to establish filiation between species, were used to construct a phylogenetic tree. A bio Neigbhor-Joining phylogenetic tree (software: PAUP*4.0a147; Retention Index = 0.49) has been built. According to the near-consensus position held within the scientific community about the recent single origin of modern humans and the fact that a part of the matriarchal mythology may spread at the time of the Out-of-Africa, the tree was rooted in East Africa.

One important counterpoint could be raised. Over time, the original character traits of a species become diluted, disappear or are even wiped out when new character traits are incorporated into the initial genome without being inherited (e.g. following symbioses or hybridization between species). In the case of myths or oral traditions, these new features are borrowings from other versions or traditions that are added to the original account. Fortunately, the methods used by biologists to study the evolution of genes can also be applied to estimating the extent of borrowing and the role of inheritance. For instance, the Retention Index (RI) allows us to measure the amount of analogous structures created by convergent evolution (homoplasy), but also measures how well the shared derived trait states (synapomorphies) explain the tree. This tool (among others) enables measurement of the percentage of traits shared by two or more tales or traditions that are not inherited from a common ancestor, i.e. to distinguish between a version / tradition that moved wholesale and one that was reconstructed out of very simple and common features (d’Huy 2012). Here the RI shows a low yet existing vertical transmission. This low score could be explained by the nature of cultural evolution with borrowings that eradicate a great part, yet not the totality, of the phylogenetic signal.

## 3. Results and discussion

In the obtained tree (figure 1; appendix A), two major clades can be identified: the first one, the Eurasian, also includes the Arctic area and western Africa; the second one, essentially Amerindian, can be subdivided into two minor clades: the most important group includes the areas of South America and the Melanesia, the other contains only an area of North America. This tree is only a model yet its structure fits very well with what is known about first human migrations^6^. Accordingly, a strong correlation between a number of motifs and genes has been shown by Andrey Korotayev and Daria Khaltourina (2011). Conversely, a low correlations between the distributions of a set of tales-type and spatial associations among populations are also consistent with vertical processes of mythological inheritance (d’Huy 2015b; Da Silva and Tehrani 2016). Moreover, phylogenetic trees based on many versions of the same myths or on many folklores associate rather strongly with the distribution of certain genes (d’Huy 2012a, b, c, 2013a, b, 2015a, b, c). So myths considered at a general level seems to respond very slowly to changes in environmental or social conditions at a similar rate than genes and could reflect a particular history of settlement, i.e. the cultural history of most of the ancestors of a given area.

**Figure 1:**
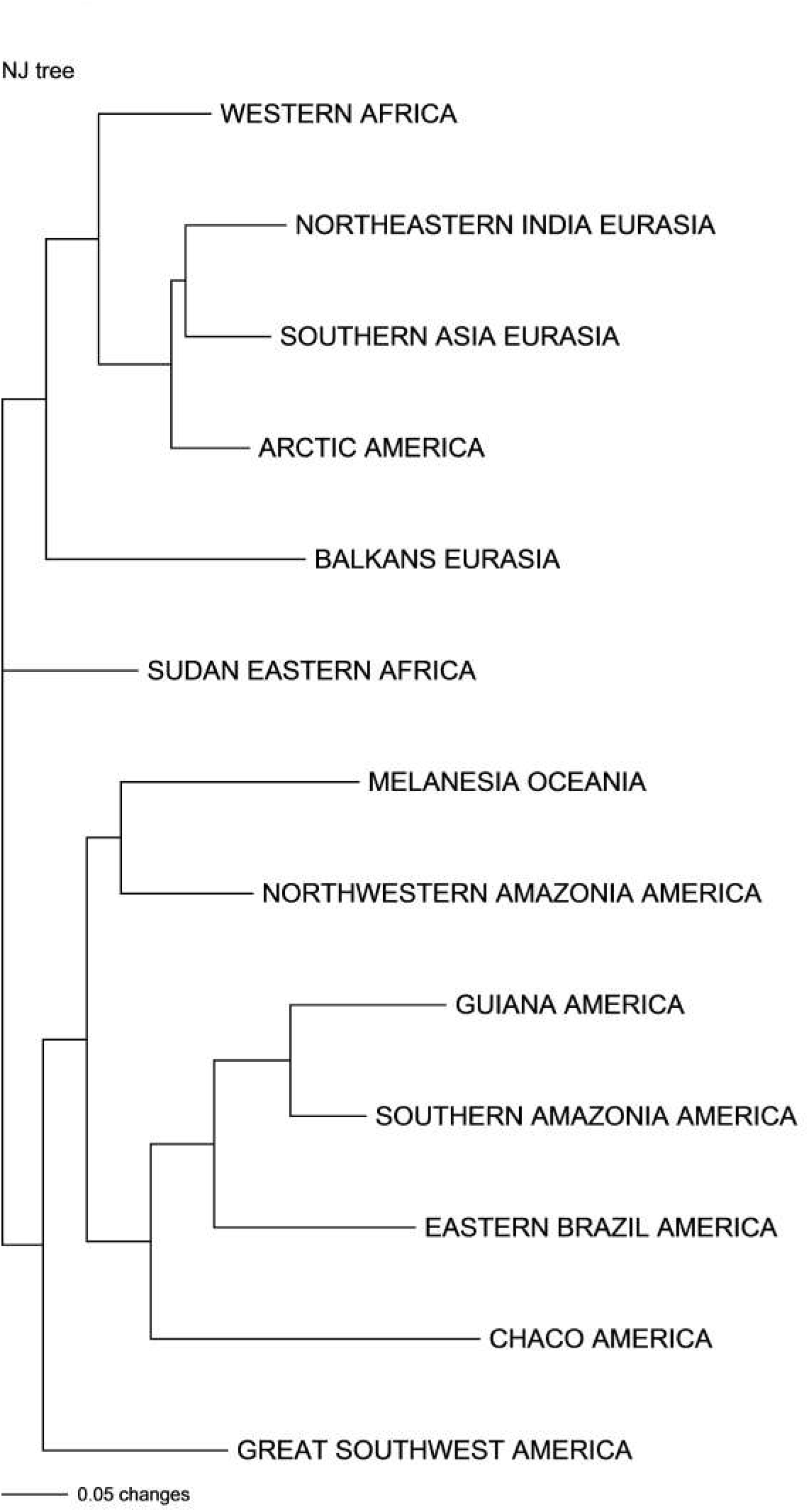
Bio Neighbor-joining tree of 13 cultural areas; narratives about the differentiations between men and women, their cohabitation and their confrontation (Berezkin’s database).

According to the tree, human beings would have gone out of Africa, between 50,000 and 100,000 years ago, followed the southern coastline of Asia and colonised Australia around 50.000 years ago (in the current tree: Melanesia). The early Amerindian people moved via the Bering Strait to South America when sea levels were significantly lowered due to the Last Glacial Maximum (in the tree: Guiana, Northwestern Amazonia, Southern Amazonia, Eastern Brazil, Chaco). The American mythology of this period is distributed throughout the Americas. The second migration from Asia, during or shortly after the initial peopling of America, has left an impact only in North America (in the tree: Great Southwest); the third migration, subsequent to the first migrations, has only left a trace in Arctic populations. The African mythology was largely lost in Continental Eurasia (in the tree: Balkans, Southern Asia, Northeastern India), likely because the decrease of population density during or soon after the Late Glacial Maximum. The place of western Africa could be explained by an early Back-to-Africa, or by an influence of the more northern regions, themselves bound to the Mediterranean culture.

It is remarkable that the structure of the tree is widely compatible with the distribution of the male rites suggested by Yuri Berezkin. It is also remarkable to find a similar tree from a completely different databases of narratives based on the motifs of dragons and serpents (d’Huy 2016a; fig. 2); the fact that results from an independent and unrelated source “converge” to the same conclusions (also see below) shows that the result of the phylogenetic method is very strong and that the output obtained by this method may be directly proportional to the input.

**Figure 2:**
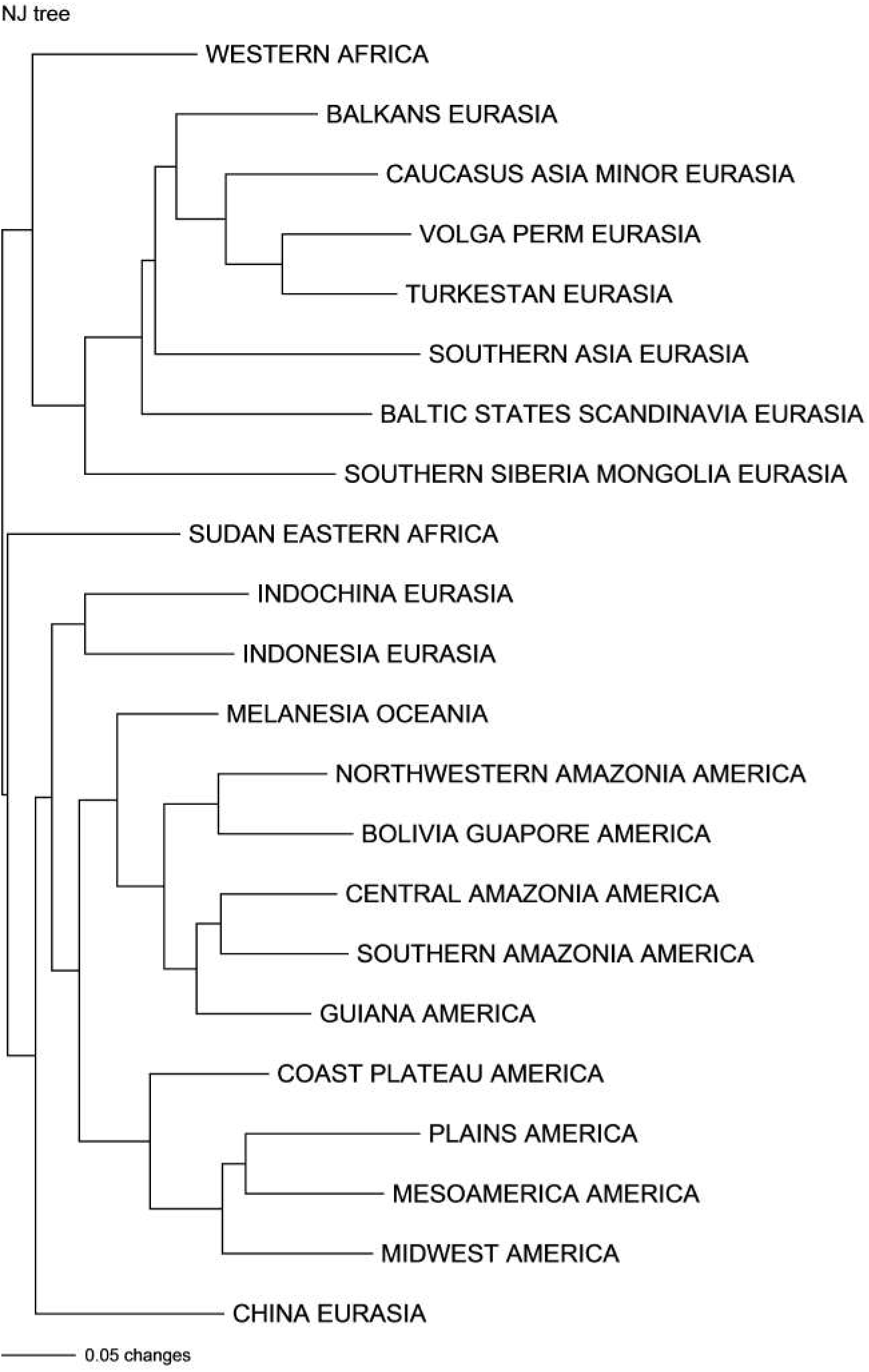
Bio Neighbor-joining tree of 22 cultural areas; narratives about serpents (Berezkin’s database).

Once the structure of the tree checked by what we know about the settlement of the planet, it becomes possible to reconstruct statistically the proto-folklore about the differentiation men/women before the Out-of-Africa. Only the features reconstructed both at the root of the tree and the cluster Melanesia / Northwestern Amazonia were retained. The reconstructed features are: F38. Women were possessors of the sacred knowledge, sanctuaries or ritual objects which are now taboo for them or they made attempts to acquire such a knowledge or objects; F40A. An anthropomorphic male or an androgyne is the only possessor or leader of women; F40B. A man gets into the village of women. Usually he has to satisfy every woman against his will or every woman claims him for herself; F42. The men feel themselves offended by the women and abandon them; F43. Women of the ancestral community kill or abandon the men; F44. Women and men quarrel and abandon each other; F45. There are (or were) women who live apart from men in their own village or villages.

In order to control the reconstructions, the results could be compared to those obtained by Andrey Korotayev and Daria Khaltourina (2011). Using principal component analysis, these researchers show the existence of a mythological cluster associating the Melanesia and the Amazonia areas. This group is believed to derive from the first waves of human migration from Africa along the southern edge of Asia and to correspond to the first colonisation of Oceania and America. Significantly, only one reconstructed feature (F40B) is not found in Korotayev and Khaltourina’s reconstructions.

|  | F16 | F38 | F39 | F40A | F40B | F41 | F42 | F43 | F44 | F45 |
| --- | --- | --- | --- | --- | --- | --- | --- | --- | --- | --- |
| Current paper |  | X | Not covered | X | X |  | X | X | X | x |
| Korotayev and Khaltourina | X | X | X | X |  | X | X | X | X | X |

Additionally, according to Yuri Berezkin, the distribution of the motif F38 and F40B at the world level fits with an African origin: the Russian researcher supposes a diffusion by the first human migrations in Australia and in New Guinea before reaching the New World via the Bering Strait (Berezkin 2013: 147–148). The phylogenetic reconstruction is thus consolidated, at least partially, by the results from two others methods^7^.

To control the existence of a “glue” holding the reconstructed folktales together since the out-of-Africa process, a new tree based only on these traits has been created. The general structure of this new tree (figure 3; appendix B) is very similar to the first one (with the exception of the Chaco area) and shows a hight Retention Index (0.7) which suggests a cohesive transmission of a set of folktales as a core tradition. According to these results, the core matriarchal folklore seems to exhibit sufficient coherence and integration to create a recognisable history when the core and the peripheral elements are taken together. Consequently, a part of the Amazon-stories should have a common origin and followed the diffusion path. Integration of these reconstructions seems to protect the core of the tradition, even if cross-cultural borrowings can locally distort the transmission.

**Figure 3:**
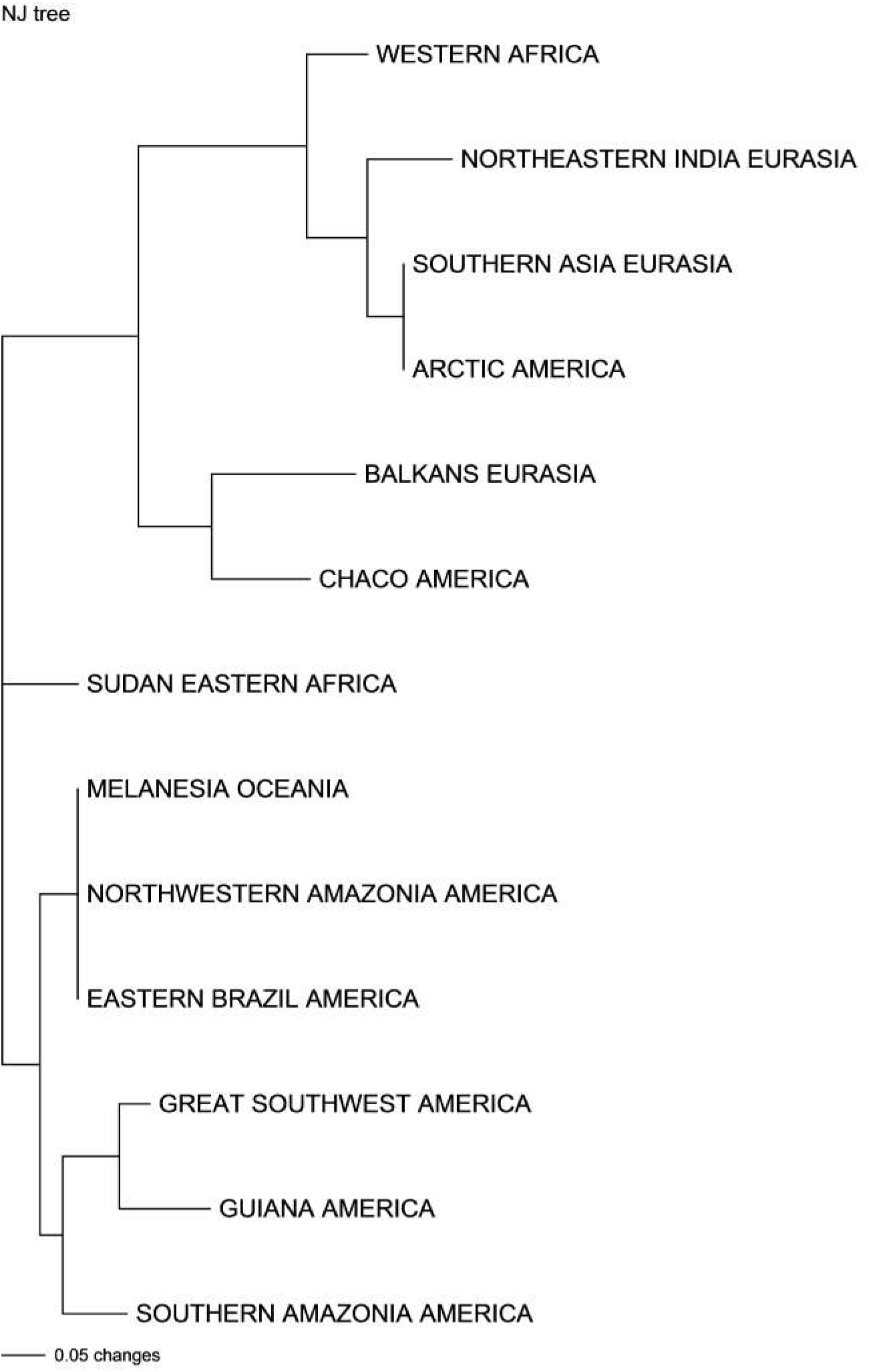
Altered bio Neighbor-joining tree of 13 cultural areas; narratives about the differentiations between men and women, their cohabitation and their confrontation.

## 4. Verification of the results

A reasonable prediction can be made about these results: if the women were supposed to dominate the men in a remote time, they also had to possess fire. To test this hypothesis, 39 other features of Berezkin’s database (D1, D1A, D1a1, D1a2, D1B, D2, D3, D4A, D4AA, D4B, D4C, D4c1, D4D, D4E, D4E1, D4F, D4G, D4H, D4h1, D4i, D4J, D4K, D4L, D4M, D4N, D4O, D4P, D5, D5A, D6A, D6B, D7, D8, D9, D10, D11, D12, D12A, D13F) concerning the acquisition of fire by the human race were used. After identifying the geographical areas possessing at least 8 narratives, a tree BioNeighborJoining (Retention Index = 0.4), rooted on East Africa, was built (fig. 4; appendix C).

**Figure 4:**
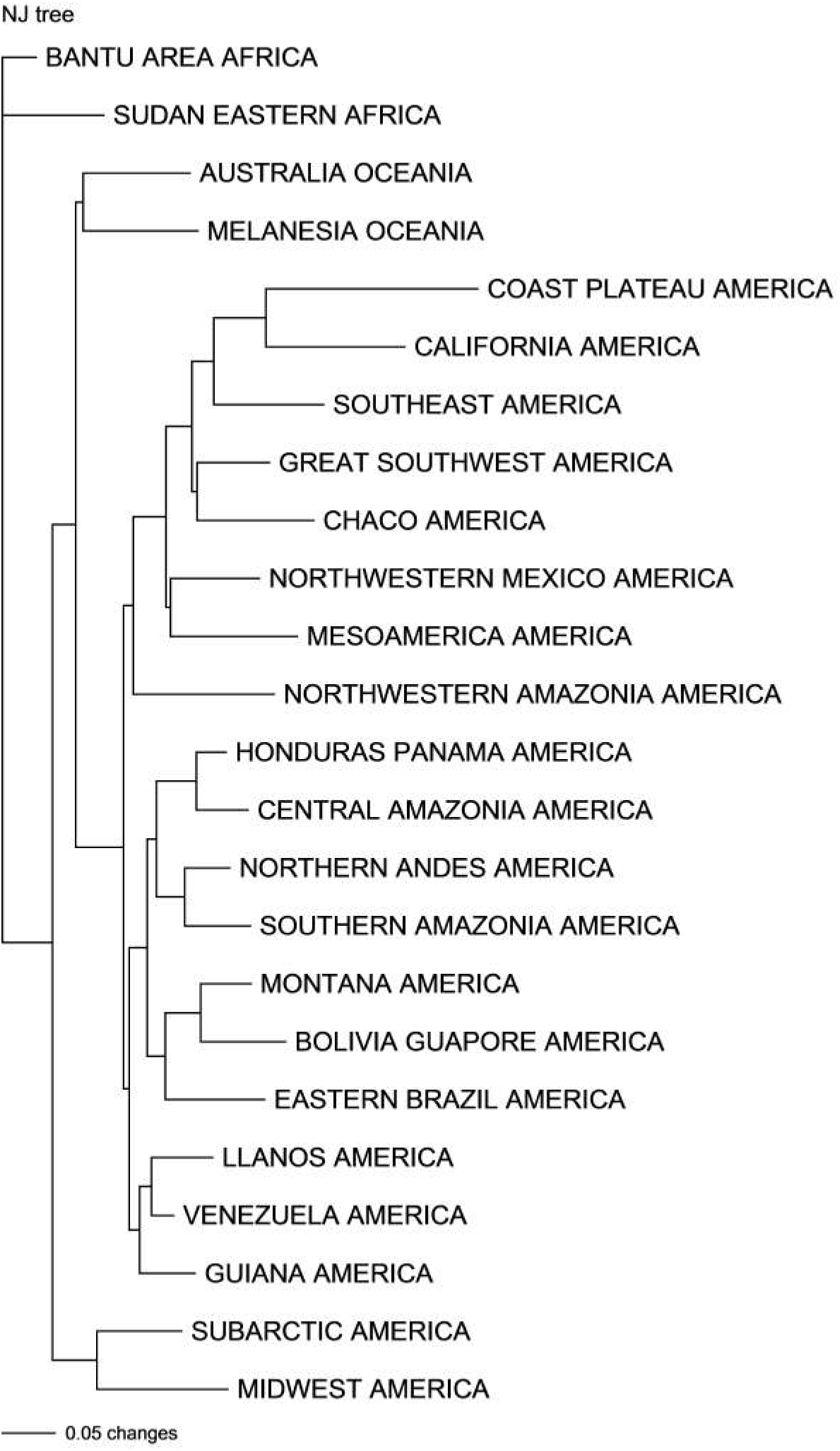
Bio Neighbor-joining tree of 24 cultural areas; narratives concerning the acquisition of fire by the human race (Berezkin’s database).

This tree presents two major amerindian clades. The first one clusters in the Midwest and the subarctic zone in North America and would represent the second expansion in North America since Eurasia (i.e. fig. 1, the Great Southwest; fig. 2, the Coast Plateau, Midwest, Great Plains and Mesoamerica) or the third expansion (i.e. fig. 1, Arctic). The second cluster, rooted in Australia and Melanesia, groups all other versions. This clade is divided into two sections: a set of areas belonging to the south of North America, Central America and the north of South America on one hand (with the exception of the Coast Plateau), and a set of South American areas on the other hand. This cluster is probably the residual trace of the first expansions in America. The place of the North-West Amazonia at the root of the first subtree suggests that there was reverse flow across the Panama isthmus after the initial settlement of South America (also see Reich et al. 2012). The Coast Plateau, the Mesoamerica and the Great Southwest areas have a rich and complex history, lying on the route of both waves of settlement, between the influence of North Amerindian and South Amerindian folklores. That explains their changing place in figures 1, 2 and 4.

The features reconstructed at the level of both the root of the tree and the node clustering Australia and Melanesia are: D4A. Fire is stolen from its original owner or brought back to the people from the person who had stolen it before; D4L. 1) First fire is sent to earth from the sky. 2) Ancestors come from the sky and bring fire or warmth from there; D5. A female personage is the owner or inventor (but not personification) of fire; D9. A raven or other big dark-feathered scavenger bird is the owner, personification, spouse, obtainer or stealer of fire, daylight, or the Sun; D12. 1) First people or inhabitants of a distant country cook food in the sun. 2) Fire owner lies that he or she cooks food in the sun.

The reconstruction of the feature D5 shows the prediction capacities of the phylogenetic method. It explains the widespread idea according to which women possessed fire before men and/or that fire springs from the body of a woman. Besides, if Frazer’s analogy between the use of the fire plough (Collina-Girard 1998) and the relationships between the sexes is accepted (according to a primitive link between the masculine stick and the feminine support) (Frazer 1930 : 220–221) this reconstruction would tend to prove that such a firelighting tool was know before the Out-of-Africa process. Note that it is nonetheless tricky to precisely determine the meaning of these primitive stories, which much depends on the context, including the social and historical context. However, the fact that the feminine power was again relegated to the remote past shows that the matriarchal myth would have been known before the Out-of-Africa process.

Frazer’s large collection of fire myths (1930; completed by Schmidt 2013) was used to create a new database to check roughly the plausibility of the previous results about the acquisition of fire. 36 features were taken into account for 9 geographic areas (with a number of features greater than 8; see appendix D) and a tree BioNeighborJoining (fig. 5; Retention Index = 0.34), rooted on Africa, was built. The amerindian clade is probably the result of the first expansions in America. The features reconstructed at the level of both the root of the tree and the node clustering Oceania, Asia and America, are: 1/ Fire in possession of women before men; 2/ Fire derived from lightning; 3/ First fire came from the sky / heaven; 4/ Fire is stolen by an animal; 5/ Fire is stolen from an animal; 6/ Tree set on fire by lightning or fire in tree.

**Figure 5:**
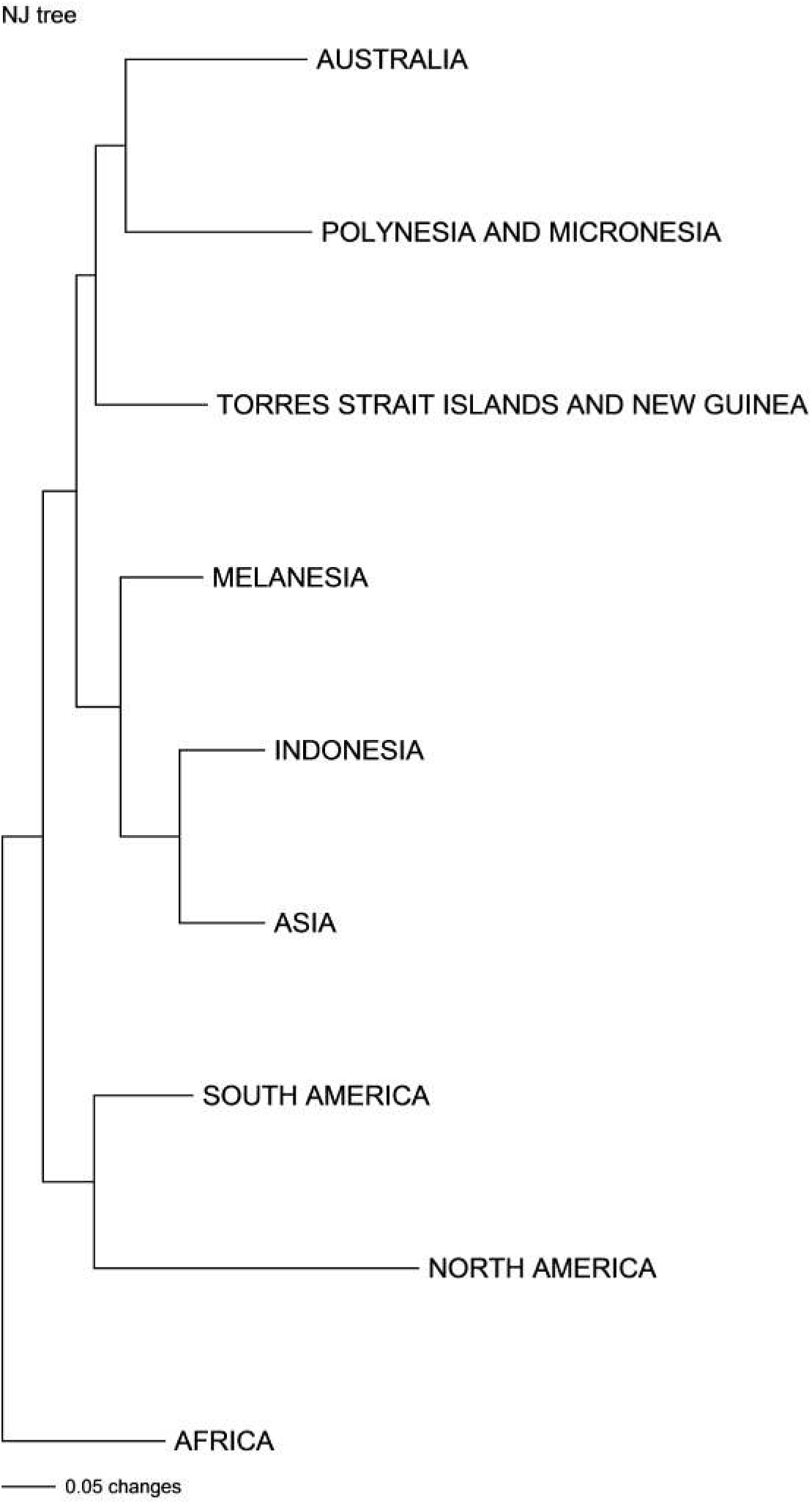
Bio Neighbor-joining tree of 9 cultural areas; narratives concerning the acquisition of fire by the human race (Frazer’s database).

In order to control the reconstructions, the two results could be compared. Only one reconstructed feature (D9) is not found in Frazer’s reconstructions.

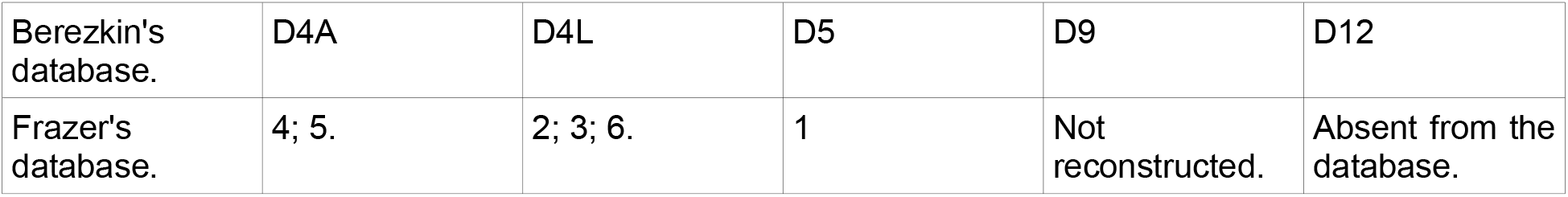

|  |  |  |  |  |  |
| --- | --- | --- | --- | --- | --- |
| Berezkin's database. | D4A | D4L | D5 | D9 | D12 |
| Frazer's database. | 4; 5. | 2; 3; 6. | 1 | Not reconstructed. | Absent from the database. |

## 5. Conclusion

These results can be harmoniously integrated into the model constructed from the trees previously reconstructed with the phylogenetic method. Indeed, taken together, mythological trees show the African origins of modern humans, who followed the southern coastline of Asia, colonised Australia around 50,000 years ago and reached America from an East Asian source. A second migration again reached North America at more or less the same time from a North Eurasian source. An admixture between the two mythological complexes took place in Siberia and in North America. However, during the Last Glacial Maximum in Eurasia, the two complexes mostly evolved in isolation, separated by natural barriers. After the Last Glacial Maximum, new stories emerged in Eurasia and spread across the continent, blurring boundaries between the primitive areas. Finally, a last flow reached America, limited to the circumpolar area, or migrated back to Asia, bringing Native American folklore (for the Out-of-Africa diffusion, see d’Huy 2013c, 2014, 2016a, b, d; for the second flow in America, see d’Huy 2012a, b; 2013a, b; 2015b; 2016c; 2017; for the third and last flow in America: d’Huy 2016c, d). This supertree of the myths would correspond to that of the genes (e.g. Raghavan et al. 2014, Skoglund et al. 2015). Additionally, several papers in comparative mythology based on different methods leads to the same conclusions (Berezkin 2007a, b, 2009, 2013; Le Quellec 2014, 2015). The fact that multiple sources of evidence are in agreement leads to very strong conclusions.

The existence of the matriarchal myth during the Out-of-Africa process indicates that the tales in which females hold primary power, predominate in roles of political leadership, moral authority, social privilege and control of property at the specific exclusion of men was already relegated to the mythical past, at least 50.000 years ago.

The author thanks Jean-Loïc Le Quellec and George Sumner for their constructive comments.

#### Box

Many reconstructed myths, using phylogenetical myths, go back to Out-of-Africa. The use of the phylogenetic method to reconstruct the past evolution of myths and folktales has been applied since 2012 (d’Huy 2012a) to numerous families of narratives^8^.

The first reconstruction of myths going back to Out-of-Africa concerned the motifs of the “dragon” and the “snake”: “*The mythological serpents guard water sources, releasing the liquid only under certain conditions. They can fly and form a rainbow. They are giant and have horns or antlers on their head. They can produce the rain and thunderstorm. Reptiles, immortal like others who shed their skin and thus rejuvenate, are contrasted with mortal men and/or are considered responsible for originating death, perhaps by their bite*.” (d’Huy 2013c, 2016a). This analysis was based on several datasets (borrowed in part from existing and already published databases such as Fontenrose 1980) and was built by varying both the definition of the “dragon” (e.g. chimera with serpentine traits; horned serpent; rainbow drinking water; snake; structural plot where a giant or lion may replace the dragon; etc) and the units of analysis (individual versions of the same tale-type, types of dragons, cultural or geographical areas) to avoid as much as possible a potential for the judgment of what stories / motifs to analyze to bias the results. Many statistical methods were also used for the same purpose. The proto-narrative, reconstructed from these various sources and analysis, probably existed prior to the Out-of-Africa process. This reconstruction was confirmed by studying the distribution of various narratives around the world and the prehistoric diffusion of these motifs (d’Huy 2016a). It was also corroborated by the comparison of many rock art images throughout the planet and by the local mythology that accompanies each of them (d’Huy 2014).

The second phylogenetic reconstruction was based exclusively on African narratives. It is a tale of the origin of death: “*The Moon sent the Hare to the people with a message of life: that men will die but be reborn as the Moon. The Hare modified the message, which then introduces death to human beings, which is why the Hare is hated by them. In punishment, the leap of the Hare is stopped, perhaps by the Moon, which explains the Hare’s lip*.” The ancient age of this narrative, and its likely presence during Out-of-Africa, was demonstrated by observing the existing patterns in the distribution of motifs (Quellec 2015a).

The third reconstruction is also exclusively based on an African corpus (Le Quellec 2015b) and the world distribution of this motif is elegantly explained by an Out-of-Africa diffusion (Berezkin 2007b; Le Quellec 2014, 2015b). The proto-tale is this: “A*nimals and men went out directly from the ground, woman came first and a part of human beings remained underground. Rock art testify today to this emergence, which can also explain the origin of “death*”.

## Footnotes

1 *“situation, dont il n’existe pas d’exemple attestés, où I’autorité est exercée exclusivement, ou principalement, par les femmes.”*

2 *“Des sociétés où le pouvoir serait entre les mains des femmes avec des hommes dominés n’existent pas et n’ont jamais existé. […] ll n’y a pas de sociétés matriarcales, parce que le modèle archaïque dominant sur toute la planète est en place dès le départ.”*

3 “*В Африке (…) встречаются аналогии тому характерному главным образом для Австралии, Меланезии и Южной Америки ритуально-мифологическому комплексу, который связан с институализированным противостоянием полов и обычно включает рассказы о былом господстве женщин не только в социальной сфере (истории о «женском царстве» в прошлом или в далекой стране известны по всему миру), но и в области культа и ритуала*.”

4 “*Можно предполагать, что мужские ритуалы, связанные с нституализированным противопоставлением полов, возникли в Африке и были принесены в встралию и на Новую Гвинею первыми сапиенсами. Через Восточную Азию этот итуально-мифологический комплекс проник в Новый Свет вместе с первой волной шедших по побережью мигрантов. (…) Из Азии полностью, а из Северной Америки в основном соответствующие мифы и ритуалы были постепенно вытеснены другими, связанными с континентально-евразийским очагом культурогенеза*.”

5 *“Myth and rituals have been misinterpreted as persistent reminders that women once had, and then lost, the seat of power. This loss accrued to them through inappropriate conduct. (…) The myths constantly reiterate that women did not know how to handle power when they had it. The loss is thereby justified so long as women choose to accept the myth. (…) In fact, if women are ever going to rule, they must rid themselves of the myth that states they have been proved unworthy of leadership roles. The final version of woman that emerges from these myths is that she represents chaos and misrule through trickery and unbridled sexuality. (…) Such visions will not bring her any closer to attaining male socioeconomic and political status, for as long as she is content to remain either goddess or child, she cannot be expected to shoulder her share of community burdens as the coequal of man. The myth of matriarchy is but the tool used to keep woman bound to her place.”*

6 A recent study has pointed out a phylogenetic signal in animal folklore that may reflect a cultural exchange between anatomically modern man and Neanderthals (d’Huy 2015b). However, such cultural exchanges between our species and other ancient hominids have not been taken into account here because of the small number of tales studied.

7 The creation of a tree also allows us to estimate the contents of certain tales at several steps in human history. For instance, it is possible to reconstruct the proto-folklore at the node of the Balkans to estimate the contents of the European Palaeolithic myths about matriarchy, i.e. F38, F40A, F40B, F44, F45 and F97 (“After eating certain fruit, berry, tuber, etc. people become sexually aware”). Such a reconstruction is of obvious interest — allowing us, for example, to rule out the hypothesis of a primitive matriarchy in interpreting the Palaeolithic Venus and representations of vulvas. Conversely, the phylogenetic reconstruction of an African myth about the emergence of humanity from the underworld (Le Quellec 2014, 2015b) or of a Palaeolithic tale about the appearance of game, delivered by a person who enters the dwelling of the master of animals (d’Huy 2013, 2015a), give an additional breadth to the interpretation of a part of this prehistoric art, seeing it as an illustration of a mythological emergence from the earth (Le Quellec 2017).

8 The worldwide distribution of these motifs is not uniform and a phylogenetic message and a geographical progression have been demonstrated, consequently the simplistic neurobiological explanation should be rejected. However, it remains possible that certain innate human reflexes, such as fear of snakes (Jones 2000, d’Huy 2013c) or a tendency to animate images of living beings (d’Huy 2013d), could contribute to a proper conservation of certain myths over time. Yet the unequal and variable distribution of the studied tales strongly implies that archetypal, neurobiological or physiological factors can not be the unique explanation of complex similarities between mythologies (d’Huy et Le Quellec 2014).

